# MNP33 is a novel mitochondrial protein that reprograms cellular bioenergetics and lipid metabolism

**DOI:** 10.1101/2025.07.29.667572

**Authors:** Yilu Xu, Quan Yuan, Yang Wang, Qiumin Liao, Xiaochuan Fu, Zhen Cao, Bin Pan, Kaixuan Zheng, Jifeng Wang, Tiemin Liu, Pingsheng Liu, Shuyan Zhang

## Abstract

Genomic regions previously annotated as non-coding are now recognized sources of functional microproteins, termed the “dark proteome”, though their identification and characterization remain challenging. Addressing this, we developed a novel proteomic strategy combining lipid droplet (LD) enrichment with mass spectrometry against the database of sORF-derived proteins. This approach yielded the first reported LD-resident microprotein, LDANP1, and subsequently revealed other candidates from the same screen, such as MNP33, were localized to mitochondria. Here, we provide the first in-depth functional characterization of MNP33, a novel 28-amino acid protein encoded by an sORF within the *Cdhr4* gene locus. We confirm MNP33 localizes to the inner mitochondrial membrane and demonstrate its interaction with the adenine nucleotide translocase 2 (ANT2). Functionally, MNP33 expression remodels mitochondrial bioenergetics, increasing basal respiration and proton leak while paradoxically elevating membrane potential, partly through modulating ANT2 activity linked to enhanced glycolysis. Furthermore, MNP33 shifts cellular lipid metabolism, favoring cholesteryl ester storage over triacylglycerols via stabilization of ACAT1, and is associated with closer ER-mitochondria contacts. MNP33 also induces ROS production and autophagy while inhibiting cell proliferation without increasing apoptosis. This study establishes MNP33 as a novel mitochondrial regulator emerging from the dark proteome, providing a mechanistic link between non-coding genetic elements, ANT2 function, mitochondrial bioenergetics, cellular metabolism, and stress responses. Our work highlights functional importances of the dark proteins and validates organelle-focused proteomics for discovering key cellular regulators.

## Introduction

The human genome contains roughly 20,000 annotated protein-coding genes, encoded by only about 1.5% of the total genomic sequence[1, 2]. However, high-throughput sequencing technologies have revealed that the eukaryotic genome is extensively transcribed, with large-scale projects like GENCODE and FANTOM indicating that at least 80% of mammalian DNA is transcriptionally active[2–6]. This has led to the identification of numerous nonclassical protein-coding genes originating from regions previously classified as noncoding[6–8]. It is now clear that many such regions harbor short open reading frames (sORFs, typically <300 nucleotides) that encode small proteins, often called microproteins or micropeptides. These sORFs are predicted to be pervasive, yet conventional gene-finding algorithms often filter them out based on size thresholds, historically overlooking a potentially significant portion of the proteins – the “dark proteins”[9–11].

Detecting these dark proteins presents challenges due to their small size, often low abundance, and potential instability[8]. Since the development of ribosome profiling provided direct evidence of translation from unexpected genomic regions, various other techniques—including advanced proteomics, HLA immunopeptidomics, CRISPR/Cas9 functional screens, computational tools and structural prediction—have been applied to systematically identify and characterize these novel proteins[11–13].

Thousands of potential non-canonical ORFs have now been predicted across genomes[11, 14]. Among these candidates, proteomic evidence, in the form of identified peptide fragments, now exists for the translation of over 1,700 non-canonical ORFs based on analysis of hundreds of datasets[15]. Furthermore, CRISPR/Cas9 screens assessing cellular phenotypes upon knockout have suggested potential functional importance for over 1,000 such microproteins[13]. For a smaller subset, currently around 15 of these proteins, the combined evidence was considered strong enough to potentially warrant their addition to official lists of protein-coding genes[15].

Importantly, dozens of human microproteins now have experimentally assigned functions, revealing their participation in diverse cellular processes[10]. Intriguingly, functional microproteins derived from noncoding sequences often localize to specific cellular membranes[16]. For instance, microproteins have been found to localize to various organelles such as the endoplasmic reticulum (ER), mitochondria, and others[17, 18]. A well-characterized group of microproteins known as “regulins”, including ELN, DWORF, MLN, ALN, PLB, and SLN, localize specifically to the ER, where they regulate the activity of the sarco/endoplasmic reticulum calcium ATPase (SERCA)[19]. These microproteins colocalize with SERCA on the ER/SR membrane, influencing calcium homeostasis and calcium-dependent signaling pathways. Additionally, several mitochondrial microproteins—such as MOTS-c, BRAWNIN, and Mitoregulin—have been identified and shown to participate in mitochondrial function and energetics[20]. Although their precise roles have yet to be fully elucidated, these novel proteins play significant roles in human physiology and diseases, including metabolism, immunology, aging, neurodegenerative disease, and cancer[11, 15, 16, 20].

Lipid droplets (LDs) are highly dynamic intracellular organelles, essential for lipid storage and metabolism, characterized by a neutral lipid core surrounded by a unique phospholipid monolayer[21, 22]. LDs are conserved from bacteria to humans[22, 23]. Despite their importance, microproteins originating from noncoding RNAs had not been previously identified as components of the LD.

To explore the potential presence of sORF-encoded microproteins in this unique organelle, we previously developed and applied an integrated strategy combining high-quality LD isolation from myoblasts with enrichment techniques for small proteins and mass spectrometry analysis against a database with sORF-derived proteins. This targeted approach successfully identified a cohort of 44 candidate microproteins derived from nominally noncoding transcripts associated with the LD fraction[24, 25].

Subsequent characterization efforts focused on validating these candidates. This led us to identify and functionally characterize LDANP1, representing the first sORF-derived microprotein demonstrated to reside on LDs[24]. Interestingly, further investigation of other candidates from this same initial cohort, published separately, revealed that several localized not to LDs but were instead found in mitochondria; among these was the protein originally designated No. 33[25].

This study provides the first in-depth functional characterization of one of these previously identified mitochondrial proteins, MNP33 (formerly No. 33), which originates from a unique sORF within the *Cdhr4* gene locus. We demonstrate that MNP33 resides in the inner mitochondrial membrane, interacts with ANT2, and significantly impacts mitochondrial bioenergetics and cellular metabolism. Our findings establish MNP33 as a novel regulatory protein emerging from the dark proteins that links noncoding elements to mitochondrial function.

## Materials and methods

### Materials

Hoechst 33342, ER Tracker Blue, LipidTOX Red, LipidTOX Deep Red and puromycin were from Invitrogen. MitoTracker Red CMXRos and PageRuler Prestained Protein Ladder were from ThermoFisher SCIENTIFIC. The ROS dye dihydroethidium (DHE) was from Beyotime. Polybrene and sodium oleate were obtained from Sigma-Aldrich. [^3^H]-oleate ([^3^H]-OA) and enhanced chemiluminescence substrate were from PerkinElmer. Polyvinylidene fluoride (PVDF) membranes were obtained from Millipore Life Sciences. Triacylglycerols Assay Kit was from BioSino Bio-Technology and Science Inc. Pierce BCA Protein Assay Kit was from ThermoFisher. Glutaraldehyde, osmium tetraoxide, uranyl acetate and lead citrate were purchased from Electron Microscopy Sciences. Antibodies against GAPDH, ACAT1, Calnexin, HMGCR, ANT2, GRP75, CPS1, CPT1A, Ki67, LC3 and Caspase3 were from ABclonal. Antibody against GFP was from Santa Cruz Biotechnology. Antibody against VDAC was from Millipore. An immuno-affinity purified rabbit polyclonal antibody targeting mouse MNP33 was generated by AbMax Biotechnology Co. The antigen, CTHLGLASSIQLPSWWSMVVNPK, was synthesized and used to immunize rabbits.

### Cell culture

HeLa cells were cultured in DMEM (Macgene Biotech., China) supplemented with 10% FBS (Hyclone), 100 U/ml penicillin and 100 μg/ml streptomycin (Macgene Biotech., China) at 37°C with 5% CO_2_.

### Plasmid construction

The MNP33 ORF was amplified from exon 10 of the mouse *Cdhr4* coding sequence. It was initially cloned into the pEGFP-N1 vector using the BamHI restriction endonuclease. The resulting recombinant plasmid (MNP33-GFP cDNA) was then further amplified and subcloned into the pQCXIP vector using NotI restriction endonuclease. The following oligonucleotides were used:

MNP33 Forward: 5L-GTACCGCGGGCCCGGATGCTACGTTGGACT-3L (Plasmid: pEGFP-N1)

MNP33 Reverse: 5L-GGCGACCGGTGGATCTTTGGGGTTGACCAC-3L (Plasmid: pEGFP-N1)

MNP33 Forward: 5L-CAGGAATTGATCCGCATGCTACGTTGGACT-3L (Plasmid: pQCXIP)

MNP33 Reverse: 5L-CATGGTGGCGCGGCCTTTGGGGTTGACCAC-3L (Plasmid: pQCXIP)

### Cell line construction

HeLa cells were cultured in 6-well plates until reaching approximately 80% confluency, then infected with pQCXIP-MNP33 pseudotyped retrovirus in the presence of 8 μg/ml polybrene to enhance infection efficiency. Starting 48 h post-infection, cells were selected with DMEM containing 1 μg/ml puromycin for at least two weeks. Subsequently, cells were seeded into 96-well plates to isolate single clones.

### Confocal microscopy

HeLa cells cultured on coverslips were transfected with 0.5 µg of the constructed MNP33 plasmid, with C-terminally fused GFP. An empty plasmid (pEGFP-N1) was used as a control. 24 h after transfection, cells were stained with MitoTracker Red or LipidTOX Red for 30 min, followed by imaging using Olympus FV1200 confocal microscope (Olympus Corp, USA). Cells stably overexpressing GFP or MNP33-GFP grown on coverslips were incubated with or without OA for 12 h and then stained with MitoTracker Red, LipidTOX Deep Red, ER Tracker Blue or TMRM for 30 min. Images were acquired using Olympus FV1200 confocal microscope.

### Isolation of lipid droplets and cellular components from HeLa cells

Lipid droplets were isolated and analyzed from HeLa cells using a previously described protocol[26]. Briefly, HeLa cells stably overexpressing GFP or MNP33-GFP were treated with or without 100 µM OA for 12 h, followed by two washes with ice-cold PBS. Cells were then scraped and resuspended in cold PBS. After centrifugation at 1,000*g* for 10 min, the resulting cell pellets were resuspended in 10 ml Buffer A (20 mM Tricine, pH 7.6, 250 mM Sucrose) containing 0.2 mM PMSF and placed on ice for 15 min. Cells were then homogenized by a N_2_ bomb at 1,000 psi for 15 min on ice. The cell homogenate was centrifuged at 1,000*g* for 10 min to remove the nuclei and cellular debris, yielding the post-nuclear supernatant (PNS) fraction. A 2 ml of PNS was then centrifuged at 8,000*g* at 4°C for 10 min to isolate mitochondria. The mitochondrial fraction was washed 2–3 times with Buffer B (20 mM HEPES, 100 mM KCl, and 2 mM MgCl_2_, pH 7.4) by centrifugation at 8,000*g* for 5 min at 4°C. The above remaining PNS was centrifuged at 182,000*g* at 4°C for 1 h. The resulting pellet was total membrane (TM) fraction. Lipid droplets appeared as an upper white layer and were carefully collected into a 1.5 ml Eppendorf tube. Collected lipid droplets were washed by centrifugation at 21,380*g* for 3 min at 4°C, and the underlying solution was discarded. Lipid droplets were gently resuspended in 200 μl Buffer B. This washing step was repeated three times. The TM fraction was also washed 2–3 times with Buffer B by centrifugation at 21,380*g* for 5 min at 4°C.

Lipid extraction and protein precipitation from lipid droplets were performed using chloroform/acetone (4:1, v/v) treatment followed by centrifugation at 21,380*g* for 10 min at 4°C. The resulting protein pellet was then dissolved in 2× SDS sample buffer [100 mM Tris-HCl (pH 6.8), 4% SDS (m/v), 20% glycerol (v/v), 4% 2-mercaptoethanol and 0.04% bromophenol blue].

### Proteinase K digestion assay

Crude mitochondrial pellets were isolated by density gradient centrifugation as described above, resuspended in Buffer B, and divided into aliquots. Serial dilutions of proteinase K were added to the mitochondrial suspensions, followed by incubation at 37°C for 15 min to allow digestion. After digestion, mitochondria were pelleted by centrifugation at 8,000*g* for 10 min at 4°C, then resuspended in Buffer B and washed twice. The final mitochondrial pellet was dissolved in 2× SDS sample buffer for subsequent analysis.

### Silver staining and Western blotting

For silver staining, proteins from different fractions were dissolved in 2× SDS sample buffer and denatured for 5 min at 95°C. Proteins were separated by sodium dodecyl sulfate-polyacrylamide gel electrophoresis (SDS-PAGE). Then at room temperature, the gels were fixed in a fixative solution (water: ethanol: acetic acid, 5:4:1, v/v/v) for 30 min and then sensitized with sensitization solution containing 30% ethanol (v/v), 12.7 mM sodium thiosulfate pentahydrate, 0.83 M sodium acetate trihydrate for 30 min. The gels were then washed four times with ddH_2_O, 5 min per wash. After washing, the gels were incubated in silver staining solution (14.72 mM silver nitrate with 4.93 μM formaldehyde added immediately before use) for 20 min. This was followed by development in chromogenic solution (0.24 M sodium carbonate with 4.93 μM formaldehyde added immediately before use) until protein bands became visible. The reaction was terminated by adding stop solution containing 43.43 mM EDTA-Na_2_ once the bands were clearly developed.

For Western blotting, proteins of different fractions were separated by SDS-PAGE and transferred to 0.2 μm PVDF membrane. Membranes were blocked with 5% BSA for 1 h at room temperature and then incubated with primary antibody for 1 h at room temperature or overnight at 4°C. Membranes were washed three times with washing buffer for 5 min each, and then incubated with the secondary antibodies for 1 h at room temperature. The membrane was washed three times with washing buffer for 5 min each and then detected by enhanced chemiluminescence (ECL).

### Transmission electron microscopy (TEM) analysis

Cells were collected by trypsin digestion and were fixed in glutaraldehyde (2.5%, v/v) in 0.1 M phosphate buffer (pH 7.2) overnight at 4°C. Subsequently, the samples were post-fixed in 1% osmium tetraoxide with 1.5% potassium ferrocyanide at 4°C for 2 h. Then the samples were dehydrated in an ethanol series and then embedded in Embed 812 to be prepared as 70-nm-thick ultrathin sections. The sections were then observed with Tecnai Spirit electron microscope (FEI, Netherlands) after stained with uranyl acetate and lead citrate.

### Analysis of mitochondrial morphology and ER–mitochondrion contacts

Mitochondrial morphology and ER-mitochondrion contacts were examined using 70-nm ultrathin cell sections imaged at 9,300× magnification under TEM. A total of 51 cells were analyzed. High-throughput quantification of mitochondrial morphology parameters and ER-mitochondrion contacts was performed using a deep learning-based method (DeepContact) integrated into Amira software. The procedures for organelle segmentation and quantification followed previously published protocols[27, 28].

### Measurement of oxygen consumption rate (OCR) and extracellular acidification rate (ECAR)

OCR and ECAR were measured using the Seahorse XFe96 Extracellular Flux Analyzer (Seahorse Bioscience, USA) in conjunction with the XF Cell Mito Stress Test Kit or Glycolysis Stress Test Kit, following the manufacturer’s instructions. Briefly, cells were seeded at 10,000 cells/well in Seahorse XF-96 plates and incubated in a CO_2_-free incubator for 1 h prior to measurement. For OCR analysis, baseline oxygen consumption was recorded first, followed by sequential injection of oligomycin (1.5 μM), FCCP (0.5 μM), and rotenone/antimycin A (0.5 μM each) to assess key parameters of mitochondrial function. Finally, basal respiration, maximal respiration, non-mitochondrial oxygen consumption, proton leak, ATP production and spare respiration capacity were calculated. For ECAR analysis, after recording the baseline, glucose (10 mM), oligomycin (1 μM), and 2-deoxyglucose (50 mM) were injected sequentially to evaluate key aspects of cellular glycolysis. All data were normalized to total protein concentration.

### ATP measurement

ATP concentration was determined using an ATP Assay Kit (Beyotime, China). Briefly, cells cultured in 6-well plates were grown to approximately 80% confluency. The plates were placed on ice, and 200 μl of cell lysis buffer was added to each well. Cells were lysed by repeated pipetting, followed by centrifugation at 12,000*g* for 5 min at 4°C. The supernatant was collected, mixed with the ATP detection reagent, and the luminescence was immediately measured using a plate luminometer or multifunctional microplate reader.

### Flow cytometry analysis of mitochondrial membrane potential (**ΔΨ**m) and reactive oxygen species (ROS)

Cells were seeded at a density of 1.0× 10^6^ cells per well and cultured for 24 h. They were then stained with TMRM (for mitochondrial membrane potential) or DHE (for ROS detection) for 30 min. Following staining, cells were trypsinized and analyzed by flow cytometry.

### Immunoprecipitation (IP)

For mitochondria protein IP, the mitochondrial pellets were isolated as described above and lysed in 1 ml of IP buffer (50 mM Tris-HCl, pH 7.5, 150 mM NaCl, 1 mM EDTA, 1% Triton X-100, 10% Glycerol) containing protease inhibitor cocktail (CWBiotech, China). Lysis was performed for 30 min at 4°C. GFP-conjugated beads were pre-blocked by incubating three times with IP buffer containing 200 μg/ml BSA. After lysis, samples were centrifuged at 17,000*g* for 10 min at 4°C, and the supernatant was collected and incubated with the blocked GFP beads for 3 h at 4°C. The beads were then washed three times with IP buffer, and bound proteins were eluted by resuspension in 2× SDS sample buffer. For whole cell lysate (WCL) IP, cells were lysed directly in IP buffer, and immunoprecipitation was performed following the same procedure as described above.

### LC-MS/MS analysis and protein identification

Excised gel bands containing immunoprecipitated proteins from duplicate experiments (Figure S3) underwent standard in-gel processing including washing, reduction with DTT, alkylation with iodoacetamide, and overnight digestion with trypsin. Peptide extracts were analyzed by nanoLC-MS/MS using an Easy n-LC 1200 HPLC system coupled to an Orbitrap Exploris 480 mass spectrometer equipped with a FAIMS Pro interface (Thermo Scientific). Peptides were loaded onto a C18 trap column and separated on a 75 μm id × 25 cm C18 analytical column (Reprosil-Pur C18 AQ, 1.9 μm) using a 73-min linear gradient. The solvent A consisted of 0.1% formic acid in water solution and the solvent B consisted of 80% acetonitrile and 0.1% formic acid. The segmented gradient was 4–9% B, 3 min; 9–20% B, 22 min; 20–30% B, 20 min; 30-40% B, 15 min; 40-95% B, 3 min; 95% B, 10min at a flow rate of 300 nl/min. FAIMS was operated using compensation voltages of -45V and -65V. Data were acquired in data-dependent mode with a 2-second cycle time. MS1 scans were acquired at 60,000 resolution (m/z 350–1500), and precursor ions were isolated (1.6 m/z window) for HCD fragmentation (NCE 28%). MS2 scans were acquired at 15,000 resolution with a maximum injection time of 22 ms and dynamic exclusion set to 40 s. Raw data files were processed using Proteome Discovery (v2.4.1.15, Sequest HT) searched against the UniProt Human database (update-09/2018). Search parameters included: no enzyme specificity, 2 allowed missed cleavages, 10 ppm precursor mass tolerance, 0.02 Da fragment mass tolerance, fixed carbamidomethylation (C), and variable oxidation (M). Protein identifications were filtered to a <1% False Discovery Rate (FDR) using Percolator.

### Prediction of interactions between MNP33 and candidate proteins

AlphaFold3 was used to predict the structure of MNP33[29], while the AlphaFold-predicted structures of ANT2, GRP75, and CPS1 were downloaded from the UniProt database (https://www.uniprot.org). Molecular docking was then performed using the GRAMM web server (https://gramm.compbio.ku.edu/), where ANT2, GRP75, and CPS1 were set as receptors, and the smaller protein, MNP33, was designated as the ligand. Free docking mode was selected for the simulations[30].The most favorable docking results, ranked by binding free energy (ΔG), are shown in Figure 3F. Docking complexes were further analyzed using PDBePISA (https://www.ebi.ac.uk/msd-srv/prot_int/pistart.html) and subsequently visualized with PyMOL (version 2.4.0), which was run in a Python 3.8 environment.

### Immunofluorescent staining of Ki67

HeLa cells stably expressing GFP or MNP33-GFP cultured on coverslips were washed three times with PBS, fixed with 4% paraformaldehyde for 30 min at room temperature, and then washed three times with PBS. Then cells were permeabilized with 0.1% Triton X-100 at room temperature and incubated with 5% BSA for 60 min at 37°C, followed by immunostaining with primary antibody against Ki67 (1:100) at 4°C overnight. Cells were washed again three times with PBS and incubated with secondary antibody goat anti-rabbit IgG H&L (TRITC) (1:200) for 1 h at 37°C. The cells were further stained with Hoechst (1:1,000) for 10 min. The cells were then observed using a confocal microscope.

### Cell apoptosis assay

Cell apoptosis was evaluated using a TUNEL Kit (Beyotime, China). Briefly, cells cultured on coverslips were washed once with PBS and fixed with 4% paraformaldehyde for 30 min. After a second PBS wash, cells were permeabilized with 0.3% Triton X-100 at room temperature for 5 min. Following two additional PBS washes, the TUNEL reaction mixture was prepared and applied according to the manufacturer’s instructions. Cells were incubated at 37°C in the dark for 60 min. Finally, the slides were washed three times with PBS and imaged using an Olympus FV1200 confocal microscope.

### Fatty acid preparation and treatment

Sodium oleate was prepared using our established method[31, 32]. Briefly, sodium oleate was added to ethanol at a final concentration of 100 mM and then sonicated on ice at 70 W, with 10-second intervals on and 3-second intervals off, until the solution became a milky solution. The prepared fatty acid stock was stored at 4°C and protected from light.

### Cellular triacylglycerol quantitation

Cellular triacylglycerol was quantified using a Triacylglycerol Assay Kit (Biosino Bio-technology & Science Inc., China). Briefly, cells cultured in 6-well plates were grown to approximately 80% confluency, then treated with or without 100 µM OA for 12 h. After treatment, cells were washed twice with ice-cold PBS, and 300 μl of 1% (v/v) Triton X-100 in PBS was added to each well. Cells were collected into 1.5 ml Eppendorf tubes and sonicated three times at 50 W using a 9-second pulse on, 2-second pulse off cycle. A 10 μl aliquot of the whole-cell lysate was used for triacylglycerol quantification. After adding the chromogenic reagent mix to the sample, the reaction mixture was incubated at 37°C for 30 min, and absorbance was subsequently measured at 505 nm. Protein concentration was determined using a BCA (Bicinchoninic Acid) Protein Assay Kit (Thermo, USA), following the manufacturer’s instructions, and used for normalization.

### Thin-layer chromatography (TLC)

Cells cultured in 6-well plates were grown to approximately 80% confluency and treated with or without 100 µM OA for various durations. Following treatment, cells were washed twice with PBS, and 300 μl of 1% (v/v) Triton X-100 in PBS was added to each well. Cells were collected into 1.5 mL Eppendorf tubes and sonicated three times at 50 W using a 9-second pulse on, 2-second pulse off cycle. Total lipids were extracted from the cell lysates using a lipid extraction reagent composed of chloroform/methanol/PBS (2:1:1, v/v/v), followed by centrifugation at 20,000*g* for 10 min at 4°C. The organic (lower) phase was carefully transferred to a new Eppendorf tube and dried under a stream of nitrogen gas. Dried lipid extracts were redissolved in chloroform and spotted onto silica gel TLC plates. Neutral lipids were separated using a solvent system of hexane/diethyl ether/acetic acid (80:20:1, v/v/v) and visualized by exposure to iodine vapor. Total protein content was measured and used as an internal control for normalization.

### Measurement of radioactive fatty acid incorporation

Cells cultured in 6-well plates were grown to approximately 80% confluency, then incubated with 2 μCi/ml [^3^H]-OA in the presence of 100 μM OA for the indicated time periods. Following treatment, cells were washed twice with PBS and lysed in PBS containing 1% Triton X-100. After collection, the lysates were sonicated and directly mixed with scintillation fluid for total radioactivity measurement using a PerkinElmer scintillation counter.

For analysis of specific lipid species, total lipids were extracted and separated by TLC, followed by visualization using iodine vapor. Individual lipid bands stained with iodine were scraped from the TLC plate, dissolved in scintillation fluid, and the incorporation of [^3^H]-OA into each lipid species was quantified using a PerkinElmer scintillation counter.

### Statistical analyses

The statistical analyses were performed using Office Excel 2010 (Microsoft Corp.) and GraphPad Prism 8 (NIH, USA). Determination of significance between groups was performed using unpaired Student t-test. Values presented are means ± SEM.

## Results

### MNP33 Is a New Mitochondrial Inner-membrane Protein Coded by a Noncoding Sequence

Proteins derived from nominally noncoding transcripts have been emerged and shown to play important roles. Previously, we developed a novel system to identify potential proteins encoded by noncoding RNAs on lipid droplets (LDs), beginning with isolated LDs and employing mass spectrometry in conjunction with a specialized sORF-derived protein database (Figure 1A)[24, 25]. Briefly, after the enrichment of small proteins of isolated LDs, the proteins are subjected to mass spectrometry (MS) analysis. It is worth noting that the MS data generated is searched against not the regular mouse protein database, but a database containing predicted proteins derived from noncoding transcripts with sORF. Therefore, noncoding RNA-encoded proteins can be identified. Among the identified proteins, LDANP1 and LDANP2, have been revealed to be novel LD-localized proteins and shown to be involved in the regulation of cellular metabolism.

**Figure 1.**
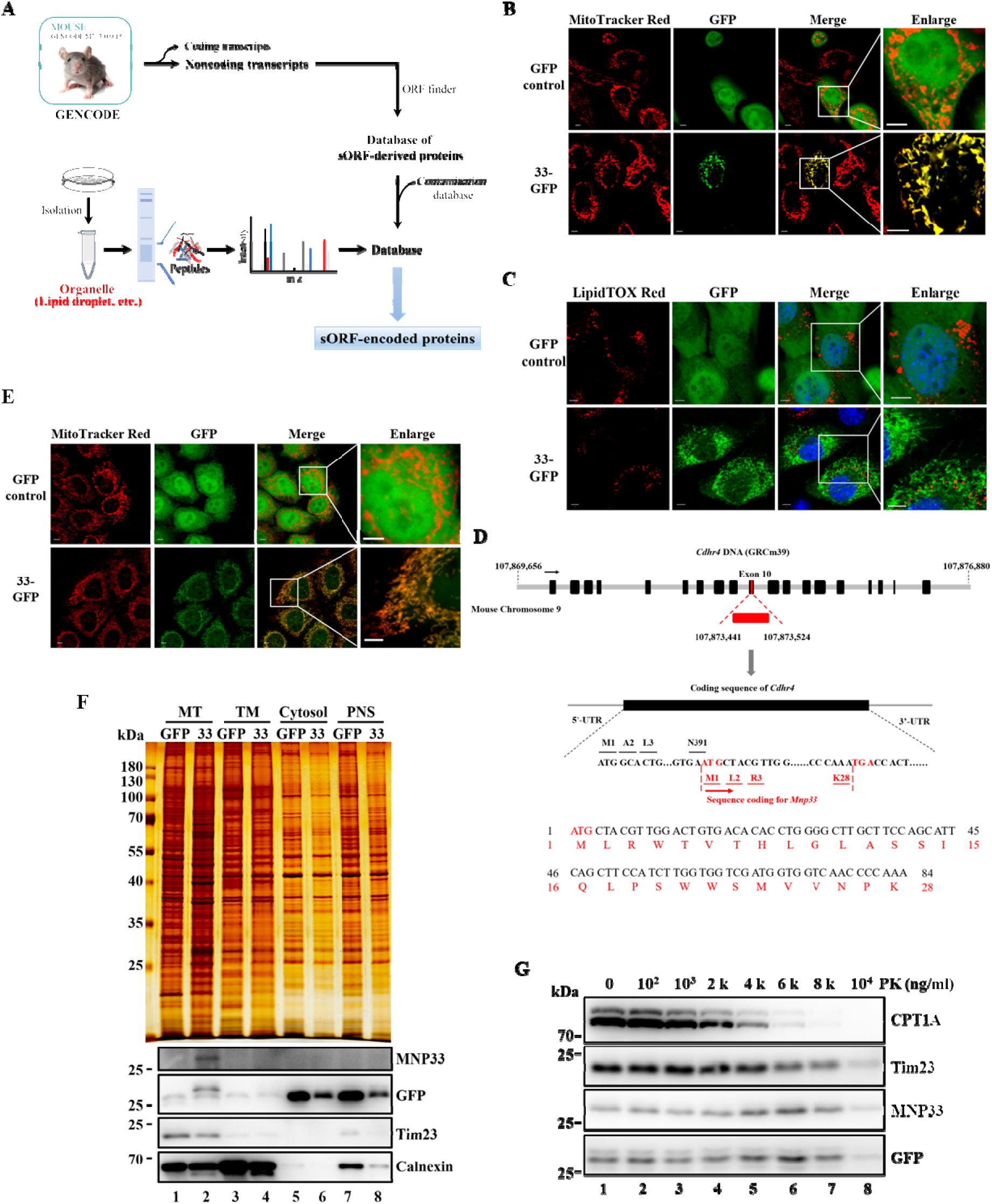
MNP33 is a Novel Mitochondrial Inner-membrane Protein. **A.** Workflow for identifying noncoding RNA-encoded proteins on LDs. The flowchart is adopted and modified from the figure in our previous work[24, 25]. **B** and **C.** Localization of transiently expressed-MNP33. GFP control or MNP33-GFP was transiently transfected into HeLa cells. Then cells were viewed using a confocal microscope after stained with MitoTracker Red for mitochondria or LipidTOX Red for LDs. Bar = 5 μm. **D.** Diagram illustrating the locus and translation of MNP33. The nucleotide and amino acid sequences of MNP33 are shown (using BioEdit) (upper panel). In mice, *MNP33* (red) is located on the exon 10 of the Cdhr4 gene (black) on chromosome 9 in Mouse GRCm39. MNP33 is translated in the +1 frame (lower panel). **E.** Localization of stably expressed-MNP33. HeLa cells stably expressing GFP or MNP33-GFP were viewed using a confocal microscope after stained with MitoTracker Red for mitochondria. Bar = 5 μm. **F.** Distribution of MNP33 assessed by subcellular fractionation. HeLa cells stably expressing GFP or MNP33-GFP were treated with 100 μM OA for 12 h and then the cells were fractionated. The proteins in different fractions were analyzed by Western blotting, probing with the antibodies indicated. 33, MNP33-GFP; MT, mitochondria; TM, total membrane; PNS, post-nuclear supernatant. **G.** Analysis of localization of MNP33 in mitochondrial membrane by proteinase K (PK) digestion assay. Mitochondria were isolated from HeLa cells stably expressing MNP33 and digested with PK at 37°C at the indicated concentrations. Western blotting analysis was performed to detect marker proteins of the mitochondrial outer membrane (CPT1A), inner membrane (Tim23), and MNP33. MNP33 was detected using antibodies against GFP or MNP33, respectively.

Interestingly, we found that another protein identified, No. 33, was shown to be localized to mitochondria[25]. Here, consistently, the transiently-expressed protein 33 with GFP tag on the C terminus shows clear GFP fluorescence overlapping with mitochondria stained by MitoTracker Red, not with LDs stained by LipidTOX Red (Figure 1B and 1C). Thus, we nominated the protein 33 as mitochondrion-associated noncoding RNA-encoded protein 33 (MNP33).

The murine *MNP33* is located to the exon 10 of cadherin-related family member 4 (*Cdhr4*) on chromosome 9, containing 84 nucleotides (from 107,873,441 to 107,873,524, in Mouse GRCm39). MNP33 is a product, containing 28 amino acids, translated in the +1 reading frame of *Cdhr4* and thus MNP33 is a novel protein, coded by a nominally noncoding sequence (Figure 1D).

To further confirm whether MNP33 is a mitochondrion-localized protein, we constructed HeLa cells stably expressing MNP33-GFP to find out its localization using both morphological and biochemical methods. The stable cells were stained with MitoTracker Red, LipidTOX Deep Red or ER Tracker Blue to mark mitochondria, LDs or ER, respectively (Figure 1E, Figure S1A and S1B). The results demonstrate that the fluorescent signals of MNP33-GFP overlap well with signals of MitoTracker Red, not with LipidTOX Deep Red or ER Tracker Blue. Thus, stably-expressed MNP33 is localized to mitochondria. Besides the morphological method, we raised a polyclonal rabbit antibody against MNP33 to further study the localization of MNP33 through analysis of its subcellular distribution after fractionation. In brief, the MNP33-GFP stable cells were fractionated after homogenization through density gradient centrifugation and the resulting fractions were analyzed by Western blotting to assess the distribution of MNP33. The results blotted with anti-GFP or anti-MNP33 antibody showed that MNP33-GFP was consistently cofractionated with known mitochondrial proteins Tim23 (Figure 1F, lane 2), but not with ER protein Calnexin (Figure 1F, lane 4). Meanwhile, MNP33 could not be detected in the LD fraction (Figure S1C, lane 2). These results demonstrate that MNP33 is a mitochondrial protein.

To determine the mitochondrial membrane localization of MNP33, mitochondria were isolated from HeLa cells stably expressing MNP33 and digested with different concentrations of proteinase K (PK) (Figure 1G). The outer mitochondrial membrane protein (OMM) CPT1A was proteolyzed at lower PK concentrations than the inner mitochondrial membrane (IMM) protein Tim23. Similar to Tim23, MNP33 was resistant to proteolysis by high concentrations of PK and exhibits a degradation pattern consistent with that of Tim23 (Figure 1G). The results indicate that MNP33 localizes to the IMM. Therefore, MNP33 is a novel mitochondrion protein localized to inner membrane of mitochondria.

### MNP33 Overexpression Alters Mitochondrial Morphology

Next, we wondered about the function of this novel mitochondrial protein. DNA sequences of MNP33 or the translated amino acid sequences of MNP33 in different species were extracted and aligned (Figure S2 and 2A). The alignment of the sequences reveals that MNP33 is highly conserved among mammals, and 60% of the amino acid residues are identical between human and mouse (Figure 2A), indicating the potential function of MNP33.

**Figure 2.**
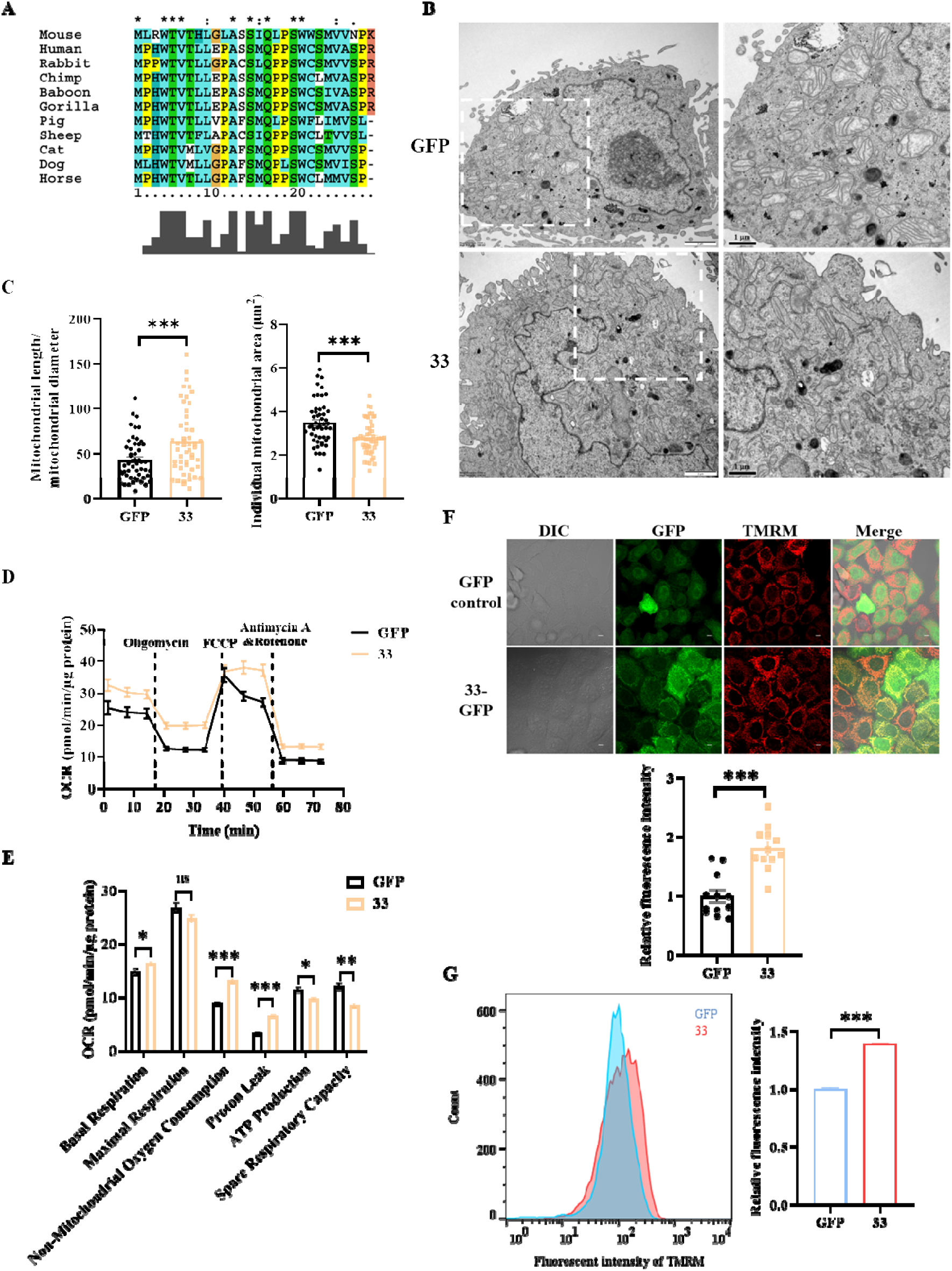
MNP33 Overexpression Induces Mitochondrial Morphological Changes and Leads to Inefficient Respiration. **A.** Conservation analysis of putative MNP33 from different mammals. The identity of the amino acid is indicated (using ClustalX). Asterisk indicates identical residues, colon represents highly similar residues, and period shows weakly similar residues. **B.** Effect of MNP33 on mitochondrial ultrastructure assessed through TEM. HeLa cells stably expressing GFP or MNP33-GFP were fixed, dehydrated, infiltrated, and embedded for ultra-thin sectioning. 70 nm sections were prepared and observed using transmission electron microscopy (TEM) to assess mitochondrial ultrastructure. Bars: 2 μm in the main image; 1 μm in the enlarged image. **C.** Quantification of mitochondrial morphology. Mitochondrial morphology parameters were analyzed using a deep learning-based approach (DeepContact) integrated into Amira software, assessing approximately 50 cells at 9,300× magnification. Left panel: Quantification of mitochondrial length-to-diameter ratio. Right panel: Quantification of individual mitochondrial area. The quantitative data were analyzed by unpaired Student t-test and presented as mean ± SEM. n=51. ***, P<0.001. **D.** Assessment of mitochondrial function using Seahorse analysis. HeLa cells stably expressing GFP or MNP33-GFP were cultured to confluence in Seahorse cell culture plates. Mitochondrial respiration was evaluated by measuring the oxygen consumption rate (OCR) under basal conditions and following sequential treatments with oligomycin (1.5 μM), the uncoupler FCCP (0.5 μM), and the Complex III and I inhibitors antimycin A and rotenone (0.5 μM each). n=5. **E.** Quantification of mitochondrial function parameters in Seahorse assays. Basal respiration, maximal respiration, non-mitochondrial oxygen consumption, proton leak, ATP production, and spare respiratory capacity were quantified from Seahorse assay data in HeLa cells stably expressing GFP or MNP33-GFP. Data were analyzed by unpaired Student t-test and presented as mean ± SEM. n=5. *, P<0.05; **, P<0.01; ***, P<0.001; ns, no significance. **F.** Observation of mitochondrial membrane potential (ΔΨm) by TMRM staining. HeLa cells stably expressing GFP and MNP33-GFP were viewed using a confocal microscope after stained with 100 nM TMRM for 30 min. Data were analyzed by unpaired Student t-test and presented as mean ± SEM. n=12. ***, P<0.001. **G.** Analysis of mitochondrial membrane potential by flow cytometry after TMRM staining. HeLa cells stably expressing GFP or MNP33-GFP were cultured to confluence, stained with TMRM for 30 min, and then analyzed by flow cytometry. Data were analyzed by unpaired Student t-test and presented as mean ± SEM. n=3. ***, P<0.001. 33, MNP33-GFP.

MNP33 was identified as a mitochondrial inner membrane protein; therefore, we first investigated its impact on mitochondrial structure. Transmission electron microscopy (TEM) revealed striking alterations in mitochondrial morphology in HeLa cells stably overexpressing MNP33. Specifically, MNP33-expressing cells exhibited significantly longer and thinner mitochondria compared to control cells (Figure 2B). Quantitative analysis of mitochondrial morphology using the DeepContact model implemented in Amira software confirmed a ∼50% increase in the length-to-diameter ratio and a decrease in the size of individual mitochondrion in MNP33-overexpressing cells (Figure 2C).

### MNP33 Increases Mitochondrial Membrane Potential Despite Enhanced Proton Leak and Inefficient ATP Production

Given the established link between mitochondrial morphology and function, we next assessed mitochondrial bioenergetics using the Seahorse XF Analyzer. MNP33 overexpression significantly enhanced basal mitochondrial respiration but did not affect maximal respiration (Figure 2D and 2E). The results also revealed that MNP33 significantly increased non-mitochondrial oxygen consumption and proton leak while reducing ATP production and spare respiratory capacity. Despite the increased proton leak, which would typically reduce mitochondrial membrane potential (ΔΨm), TMRM staining indicated an increase in ΔΨm (Figure 2F and 2G). This suggests that other factors are significantly contributing to the mitochondrial membrane potential in MNP33 cells, outweighing the effect of the leak.

Collectively, these results suggest that MNP33 induces a more “wasteful” mitochondrial state, characterized by higher oxygen consumption, increased membrane potential, but inefficient ATP production.

### MNP33 Enhances Glycolysis and Increases Total Cellular ATP

Surprisingly, despite the reduced efficiency of mitochondrial ATP production, total cellular ATP levels were increased in MNP33-overexpressing cells, as determined by ATP assays (Figure 3A). To understand this discrepancy, we measured the extracellular acidification rate (ECAR) using the Seahorse XF Analyzer as an indicator of glycolysis. MNP33 overexpression significantly increased basal glycolysis and decreased glycolytic reserve, although glycolytic capacity remained unchanged (Figure 3B and 3C). These results imply that MNP33 shifts cells to rely more on glycolysis for ATP generation, compensating for reduced ATP production via OXPHOS and netting a rise in overall ATP levels.

### ANT2 Is Identified as a Potential MNP33-interacting Protein

Based on its amino acid sequence, MNP33 appears to lack any recognizable enzymatic motifs. In addition, small proteins derived from nominally noncoding RNA sequences often exert their function through interactions with other proteins[8]. To identify potential protein interactors that could shed light on the role of MNP33 in modulating mitochondrial membrane potential and cellular bioenergetics, we sought to identify its interacting proteins.

To identify potential MNP33 interacting proteins, mitochondria were isolated from HeLa cells stably expressing MNP33-GFP, and co-immunoprecipitation (Co-IP) was performed using anti-GFP beads. Silver staining of the immunoprecipitates revealed distinct protein bands (designated Bands 1, 2, and 3) that were uniquely present or highly enriched in the MNP33-GFP IP sample compared to controls (Figure 3D, lane 3 vs lanes 1 and 2). Western blotting using an anti-GFP antibody confirmed that the band just below Band 3 corresponded to the immunoprecipitated MNP33-GFP protein itself (Figure 3E, lane 3). The Co-IP experiments were conducted in duplicate, yielding a consistent pattern of these unique bands across replicates (Figure S3). These three specific bands (Bands 1, 2, and 3) were excised from the gel and subjected to mass spectrometry analysis (Table S1). The identified proteins with ≥3 unique peptides were further analyzed. Notably, the analysis identified CPS1, GRP75, and ANT2 as the top-ranked protein hits (Top 1) for Bands 1, 2, and 3, respectively, consistent across both independent biological replicates (Table S2). This top ranking was supported by high-confidence metrics including molecular weight correspondence, unique peptide counts, and protein scores. Therefore, these consistently top-ranked candidates—CPS1, GRP75, and ANT2—were selected for further validation as potential MNP33 interactors.

To validate these interactions, Western blotting confirmed the presence of CPS1, GRP75, and ANT2 in the MNP33-GFP IP samples (Figure 3E, lane 3). ANT2 exhibited the highest enrichment ratio compared to the input (Figure 3E, lane 3 compared to lane 6), indicating a stronger interaction with MNP33 than CPS1 and GRP75. Protein-protein docking analysis using GRAMM suggests that MNP33 had the lowest predicted ΔG when interacting with ANT2 compared to CPS1 or GRP75 (Figure 3F), indicating a more energetically favorable interaction. A hypothetical model of the MNP33-ANT2 interaction generated using PyMOL is shown in Figure 3G. However, it is important to note that this model requires experimental validation.

Collectively, these results indicate that ANT2 could be the most prominent interactor with MNP33, likely underpinning the observed increases in mitochondrial membrane potential and altered bioenergetics mediated by MNP33.

### MNP33 Modulates ANT2 Activity to Maintain Elevated Mitochondrial Membrane Potential

Further we investigated how MNP33 maintains its elevated mitochondrial membrane potential (ΔΨm) despite increased proton leak and inefficient respiration, focusing on its interaction with ANT2, as ANT2 activity has been reported to contribute to the magnitude of ΔΨm by importing cytosolic ATP^4-^ in exchange for matrix ADP^3-^, resulting in a net import of negative charge[33]. Baseline TMRM fluorescence confirmed a higher ΔΨm in MNP33-overexpressing cells (Figure 3H). In control cells, the Complex I inhibitor rotenone decreased ΔΨm, and subsequent addition of the ANT inhibitor atractyloside had no further effect, indicating minimal ANT dependence when the electron transport chain is compromised (Figure 3H). However, in MNP33-overexpressing cells, while rotenone also decreased ΔΨm (still remaining higher than controls with rotenone), subsequent addition of atractyloside caused a further significant decrease in ΔΨm (Figure 3H). This indicates that, particularly when oxidative phosphorylation is impaired, ANT2 activity plays a crucial role in maintaining the elevated ΔΨm in MNP33 cells, likely fueled by the enhanced glycolysis and high cytosolic ATP levels in these cells. Indeed, inhibiting glycolysis with 2-deoxyglucose (2-DG), a glucose analog, significantly reduced ΔΨm in MNP33 cells but not in controls (Figure 3I), further highlighting the reliance of this elevated potential on glycolytically-fueled ATP import via ANT2. MNP33 overexpression did not change ANT2 expression levels (Figure 3J).

**Figure 3.**
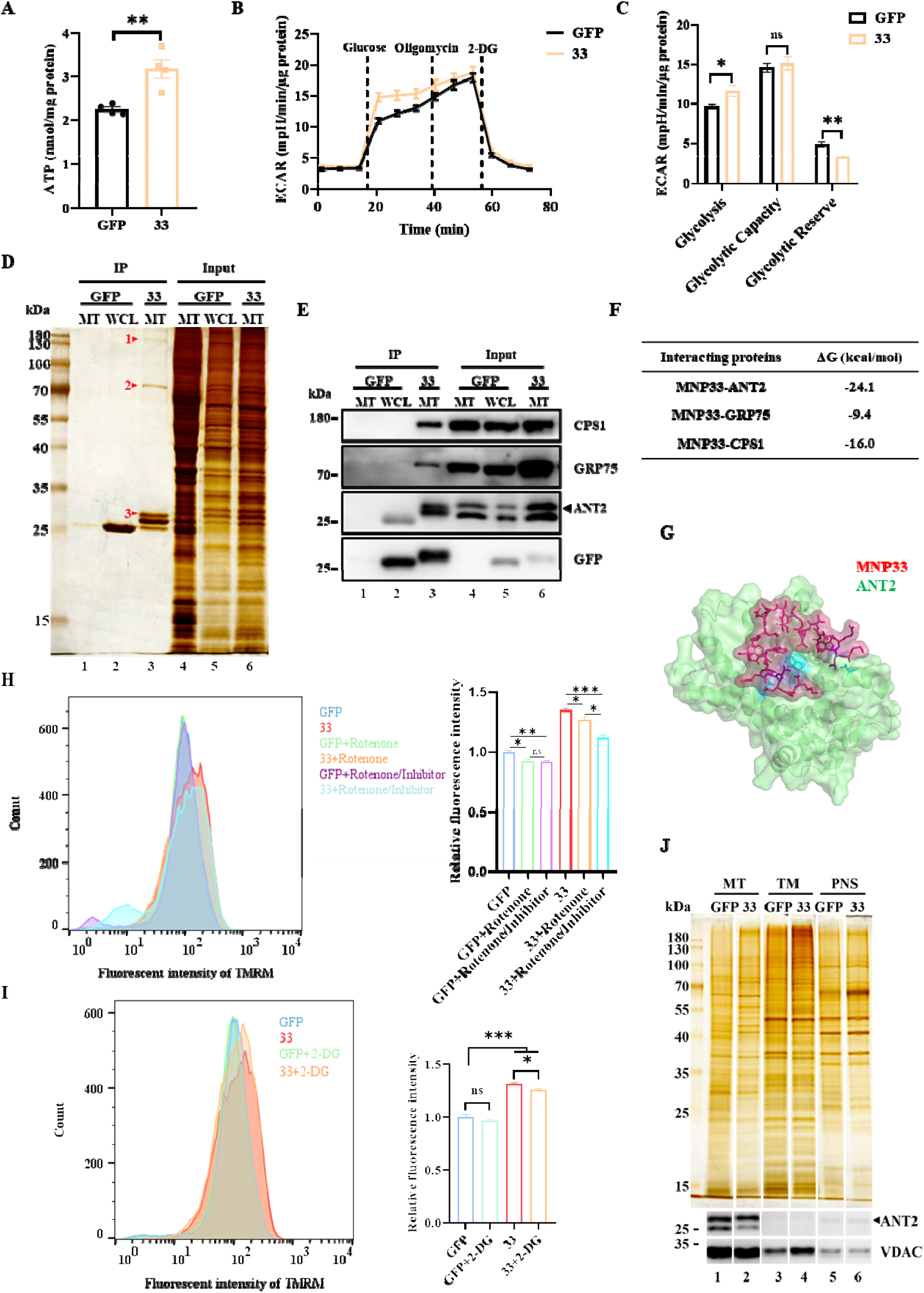
MNP33 Promotes Glycolysis and Elevates Cellular ATP Levels, Potentially Modulates ANT2 Activity to Enhance Mitochondrial Membrane Potential. **A.** Detection of ATP content in HeLa cells stably expressing GFP or MNP33-GFP. HeLa cells stably expressing GFP or MNP33-GFP were grown to confluence and ATP concentration was measured using an ATP detection kit. Data were analyzed by unpaired Student t-test and presented as mean ± SEM. n=4. **, P<0.01. **B.** Evaluation of cellular glycolysis using Seahorse analysis. HeLa cells stably expressing GFP or MNP33-GFP were cultured to confluence in Seahorse culture plates. Glycolytic function was assessed by sequentially treating the cells with glucose (10 mM), oligomycin (1 μM), and 2-deoxyglucose (2-DG, 50 mM), while recording the extracellular acidification rate (ECAR) profiles. n=5. **C.** Quantification of cellular glycolysis parameters in Seahorse assays. Glycolysis, glycolytic capacity, and glycolytic reserve were quantified in HeLa cells stably expressing GFP or MNP33-GFP using Seahorse assay data. Data were analyzed by unpaired Student t-test and presented as mean ± SEM. n=5. *, P<0.05; **, P<0.01; ns, no significance. **D-E.** Immunoprecipitation and Western blotting analysis of MNP33-interacting proteins. HeLa cells stably expressing GFP or MNP33-GFP were collected, and mitochondria were isolated by fractionation. Immunoprecipitation with anti-GFP beads was performed on the mitochondrial fraction, followed by silver staining (**D**) and Western blotting (**E**) of the resulting immunoprecipitates. 33, MNP33-GFP; MT, mitochondria; WCL, whole cell lysate. **F.** Prediction and analysis of MNP33 binding strength with interacting proteins. The binding strength of MNP33 with each of the three interacting proteins was predicted using GRAMM. The resulting predictions were analyzed and presented using PDBePISA. **G.** Protein–protein docking of MNP33 and ANT2. Protein – protein docking visualization of MNP33 and ANT2 was performed using PyMOL. **H.** TMRM staining with rotenone and ANT inhibitor treatment. HeLa cells stably expressing GFP or MNP33-GFP were grown to confluence, then treated with 0.5 μM rotenone for 20 min, or with 0.5 μM rotenone for 20 min followed by 50 μM atractyloside, an ANT inhibitor, for 30 min. After treatment, cells were stained with TMRM and analyzed by flow cytometry. Data were analyzed by unpaired Student t-test and presented as mean ± SEM. n=3. *, P<0.05; **, P<0.01; ***, P<0.001; ns, no significance. **I.** TMRM staining following 2-DG treatment. HeLa cells stably expressing GFP or MNP33-GFP were grown to confluence, treated with or without 50 mM 2-deoxyglucose (2-DG) for 20 min, stained with TMRM, and then analyzed by flow cytometry. Data were analyzed by unpaired Student t-test and presented as mean ± SEM. n=3. *, P<0.05; ***, P<0.001; ns, no significance. **J.** Effects of MNP33 on ANT2 expression through Western blotting. HeLa cells stably expressing GFP or MNP33-GFP were collected, fractionated, and analyzed by Western blotting with the indicated antibodies to assess the impact of MNP33 on ANT2 expression. 33, MNP33-GFP; MT, mitochondria; TM, total membrane; PNS, post-nuclear supernatant.

These results suggest that MNP33 modulates ANT2 activity to sustain a higher mitochondrial membrane potential.

### MNP33 Increases ROS Production, Induces Autophagy, and Inhibits Cell Proliferation

Given the observed alterations in energy metabolism, we next examined the impact of MNP33 on cellular stress responses. MNP33 overexpression significantly increased cellular reactive oxygen species (ROS) levels, as measured by DHE staining and flow cytometry (Figure 4A). To assess the cellular consequences of increased ROS, we examined cell proliferation, autophagy, and apoptosis. MNP33 overexpression was suggested to suppress cell proliferation, as evidenced by Ki67 immunofluorescence revealing a marked reduction in actively dividing cells (Figure 4B). Western blotting analysis demonstrated a marked increase in LC3 II levels within both isolated mitochondrial and total membrane fractions, suggesting enhanced autophagy, including mitophagy (Figure 4C). Notably, levels of cleaved Caspase3, an apoptosis marker, remained unchanged. Consistently, TUNEL staining showed no significant difference in apoptotic cells between MNP33-expressing and control groups (Figure 4D).

**Figure 4.**
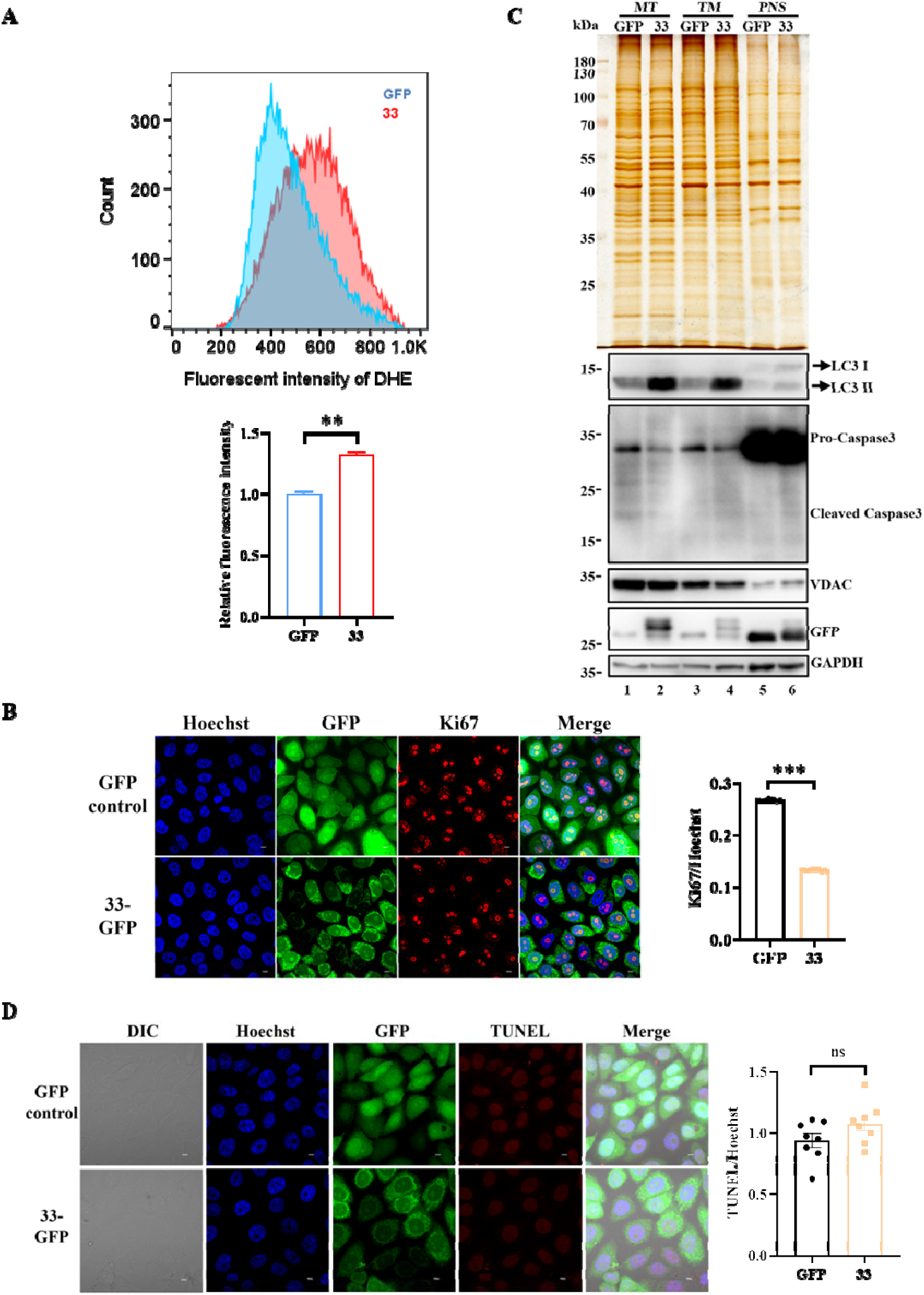
MNP33 Enhances ROS Production, Promotes Autophagy, and Suppresses Cell Proliferation. **A.** Analysis of cellular ROS by flow cytometry after DHE staining. HeLa cells stably expressing GFP or MNP33-GFP were cultured to confluence, stained with 10 μM DHE for 30 min, and subsequently analyzed by flow cytometry. Data were analyzed by unpaired Student t-test and presented as mean ± SEM. n=3. **, P<0.01. **B.** Immunofluorescence microscopy of Ki67. HeLa cells stably expressing GFP or MNP33-GFP were fixed, incubated overnight with a Ki67 antibody, then labeled with a red fluorescent secondary antibody and counterstained with Hoechst for DNA, before observation under a confocal microscope. Data were analyzed by unpaired Student t-test and presented as mean ± SEM. n=10. ***, P<0.001. **C.** Assessment of autophagy and apoptosis by Western blotting. HeLa cells stably expressing GFP or MNP33-GFP were collected, fractionated, and analyzed by Western blotting with the indicated antibodies to detect proteins involved in autophagy and apoptosis. 33, MNP33-GFP; MT, mitochondria; TM, total membrane; PNS, post-nuclear supernatant. **D.** Assessment of apoptosis by TUNEL staining. HeLa cells stably expressing GFP or MNP33-GFP were fixed and stained with TUNEL to evaluate apoptosis and then observed under confocal microscopy. Data were analyzed by unpaired Student t-test and presented as mean ± SEM. n=8. ns, no significance.

These findings suggest that MNP33-induced mitochondrial dysfunction leads to increased ROS production, triggering autophagy and inhibiting cell proliferation, but not apoptosis.

### MNP33 Reduces Triacylglycerol Accumulation and Promotes Cholesteryl Ester Storage

Considering the central role of mitochondria in lipid metabolism, we then investigated the impact of MNP33 on lipid droplets (LDs), the primary storage sites for neutral lipids. LDs were stained with LipidTOX Red and visualized by confocal microscopy after cells were treated with oleic acid (OA) for 12 h. The results showed a significant reduction in the number of LDs in MNP33-GFP cells, with LD size remained unchanged (Figure 5A). Consistently, LDs were isolated after OA treatment for 12 h, and their sizes were measured using a Delsa Nano C Particle Analyzer. The results also showed that there was no significant difference in LD size between two cell lines (Figure 5B). To further explore the effect of MNP33 on lipid metabolism, we measured triacylglycerol (TAG) levels in cells treated with or without oleate (OA). Under basal conditions, MNP33 did not affect total TAG levels. However, after prolonged OA treatment (12 h), MNP33 significantly reduced TAG accumulation (Figure 5C). Time-course experiments revealed that TAG levels were similar between control and MNP33-overexpressing cells at early time points (0, 2, 4 h) but significantly lower in MNP33 cells after extended OA treatment (8 h, 12 h) (Figure 5D). To gain a more comprehensive understanding of lipid metabolism, we analyzed another major neutral lipid, cholesteryl ester (CE), using thin-layer chromatography (TLC). Notably, CE levels were significantly increased in MNP33-overexpressing cells at all time points (Figure 5E and 5F).

**Figure 5.**
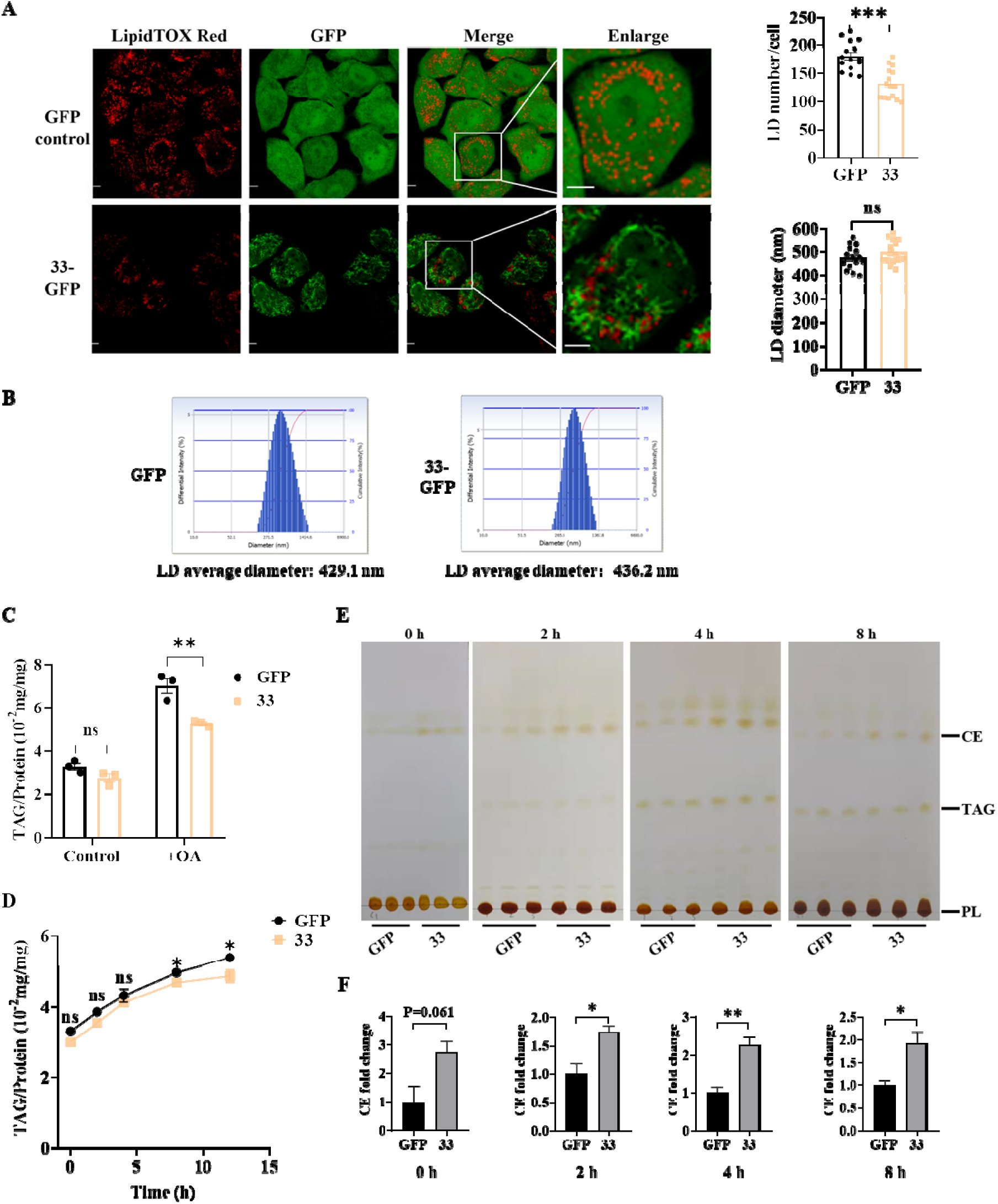
MNP33 Alters Cellular Lipid Metabolism, Favoring Cholesteryl Ester Storage Over Triacylglycerol Accumulation. **A.** Effect of MNP33 on cellular lipid droplets. HeLa cells stably overexpressing GFP or MNP33-GFP were treated with 100 μM oleic acid (OA) for 12 h, stained with LipidTOX Red to visualize lipid droplets (LDs), and then imaged via confocal microscopy. The number and size of LDs were quantified using ImageJ. Bar=5 μm. Data were analyzed by unpaired Student t-test and presented as mean ± SEM. n=15. ns, no significance. ***, P < 0.001. **B.** Size analysis of isolated LDs. HeLa cells stably overexpressing GFP or MNP33-GFP were treated with 100 μM OA for 12 h. The isolated LDs were then analyzed for size distribution using a Delsa Nano C particle analyzer. **C.** Effect of MNP33 on intracellular triacylglycerol (TAG) storage. HeLa cells stably overexpressing GFP or MNP33-GFP were cultured to confluence in 6-well plates and treated with or without 100 μM OA for 12 h. The cells were collected in PBS containing 1% (v/v) Triton X-100, sonicated, and assessed for TAG content using a commercial TAG assay kit. Data were analyzed by unpaired Student t-test and presented as mean ± SEM. n=3. ns, no significance. **, P < 0.01. **D.** Time course analysis of intracellular TAG content after OA treatment. HeLa cells stably overexpressing GFP or MNP33-GFP were treated with 100 μM OA for the indicated durations. TAG levels were subsequently measured using TAG assay kit and normalized against total protein. Data were analyzed by unpaired Student t-test and represented as mean ± SEM (n=3). ns, no significance; *, P < 0.05. **E.** Thin-layer chromatography (TLC) analysis of intracellular cholesteryl ester (CE). HeLa cells stably overexpressing GFP or MNP33-GFP were treated with 100 μM OA for the indicated times. After sonication, protein concentrations were measured, and equal amounts of total protein were used for lipid extraction. The extracted lipids were separated by TLC and visualized using iodine vapor. TAG, triacylglycerol; CE, cholesteryl ester; PL, phospholipids. **F.** Quantification of CE from TLC plates. CE levels were quantified by measuring the gray intensity of TLC bands using ImageJ. Data were analyzed by unpaired Student t-test and presented as mean ± SEM. n=3. *, P < 0.05; **, P < 0.01; ns, no significance.

These data suggest that MNP33 overexpression shifts cellular lipid metabolism, promoting CE storage over TAG accumulation.

### MNP33 Stabilizes ACAT1 and Alters ER-Mitochondrion Proximity

To elucidate the mechanisms underlying the increased CE accumulation, we first examined fatty acid uptake using radiolabeled OA ([^3^H]-OA). MNP33 significantly reduced [^3^H]-OA uptake by cells at 2 h and 8 h (Figure 6A). We then analyzed the incorporation of [^3^H]-OA into different lipid classes by separating them via TLC. The lipid spots were then scraped, and their radioactivity was measured using scintillation counting. MNP33 significantly reduced the incorporation of [^3^H]-OA into newly synthesized TAG at 2, 4, and 8 h (Figure 6B). Conversely, MNP33 significantly increased the incorporation of [^3^H]-OA into CE and phospholipids (PL) after 8 h of OA treatment (Figure 6C and 6D).

**Figure 6.**
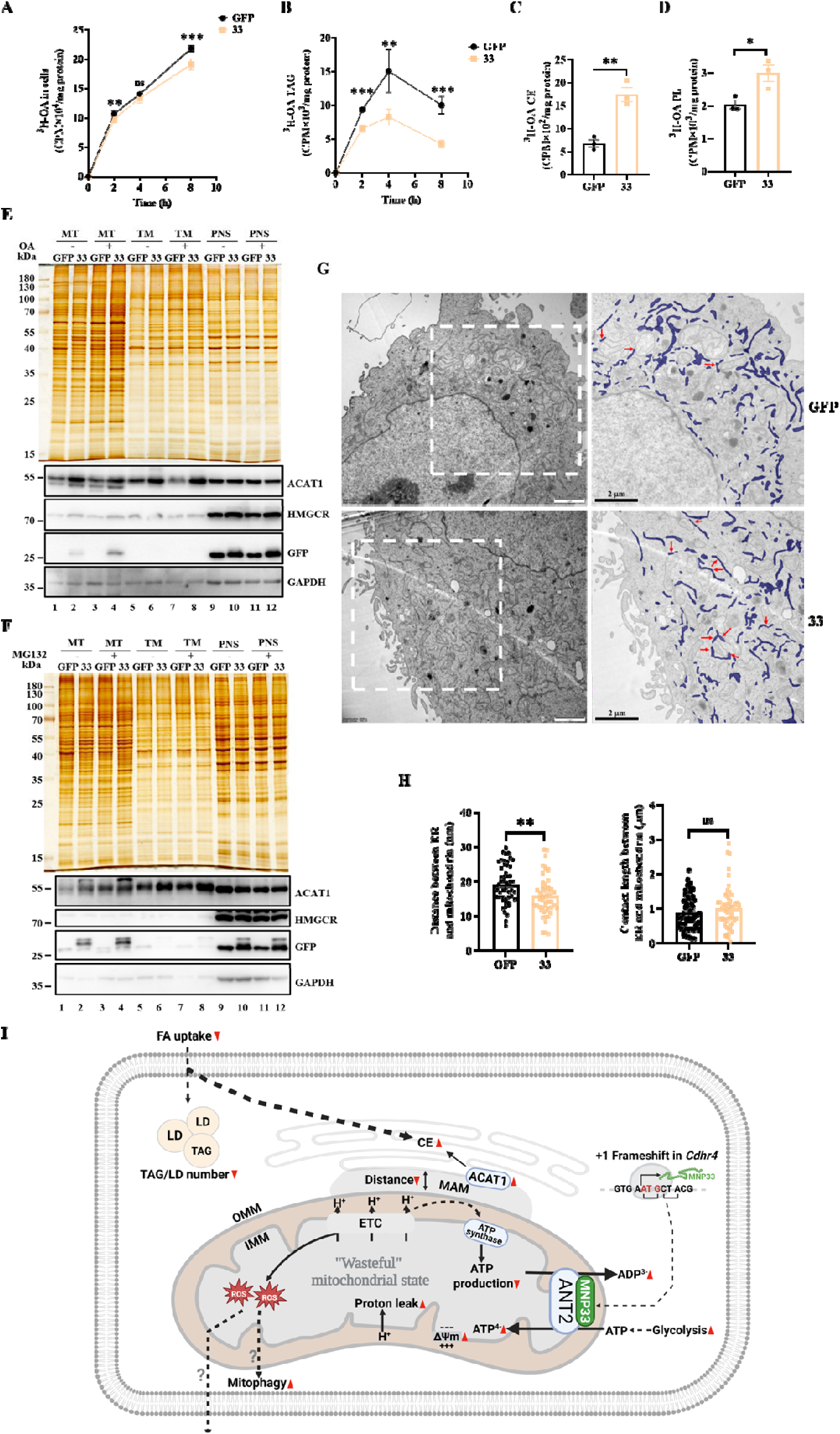
MNP33 Promotes Cholesteryl Ester Accumulation through ACAT1 Stabilization and Enhances ER-Mitochondrion Contact. **A.** Measurement of cellular uptake of [^3^H]-OA. HeLa cells stably overexpressing GFP or MNP33-GFP were incubated with 2 μCi/ml [^3^H]-OA in the presence of 100 μM OA for the times indicated. Total lipids were then extracted and uptake of [^3^H]-OA was measured by scintillation counting. Data were analyzed by unpaired Student t-test and presented as mean ± SEM. n=3. ns, no significance; **, P < 0.01; ***, P < 0.001. **B.** Measurement of [^3^H]-labeled TAG in HeLa cells stably overexpressing GFP or MNP33-GFP were incubated with 2 μCi/ml [^3^H]-OA in the presence of 100 μM OA for the various times indicated. Total lipids were then extracted and the [^3^H]-labeled TAG was separated by TLC. [^3^H]-labeled TAG was collected from TLC plate by scraping and measured by scintillation counting. Data were analyzed by unpaired Student t-test and presented as mean ± SEM. n=3. **, P < 0.01; ***, P < 0.001. **C** and **D.** Measurement of [^3^H]-labeled CE and PL. HeLa cells stably overexpressing GFP or MNP33-GFP were incubated with 2 μCi/ml [^3^H]-OA in the presence of 100 μM OA for 8 h. Total lipids were then extracted, and [^3^H]-labeled CE (**C**) and phospholipids (PL) (**D**) were separated by TLC. The corresponding lipid spots were scraped from the TLC plate and quantified using scintillation counting. Data were analyzed by unpaired Student t-test and presented as mean ± SEM. n=3. *, P < 0.05; **, P < 0.01. **E.** Western blotting analysis of key enzymes in cholesteryl ester synthesis. HeLa cells stably overexpressing GFP or MNP33-GFP were collected and fractionated following a 12-hour treatment with or without 100 μM OA. ACAT1 and HMGCR expression levels were analyzed by Western blotting, with GAPDH and silver staining used as internal controls. 33, MNP33-GFP; MT, mitochondria; TM, total membrane; PNS, post-nuclear supernatant. **F.** Effect of MNP33 on ACAT1 stability. HeLa cells stably overexpressing GFP or MNP33-GFP were collected and fractionated following a 12-hour treatment with or without 1 μM proteasome inhibitor MG132. Protein levels in different fractions were analyzed by Western blotting using the indicated antibodies, with GAPDH and silver staining serving as internal controls. 33, MNP33-GFP; MT, mitochondria; TM, total membrane; PNS, post-nuclear supernatant. **G.** Effect of MNP33 on ER-mitochondrion contacts assessed through TEM. HeLa cells stably expressing GFP or MNP33-GFP were fixed, dehydrated, infiltrated, and embedded for ultra-thin sectioning. 70 nm sections were prepared and observed using transmission electron microscopy (TEM) to assess ER-mitochondrion contacts. Bar=2 μm. **H.** Quantification of ER-mitochondrion contacts. ER-mitochondrion contacts were analyzed using a deep learning-based approach (DeepContact) integrated into Amira software, assessing approximately 50 cells at 9,300× magnification. Left panel: Quantification of distance between ER and mitochondria. Right panel: Quantification of contact length between ER and mitochondria. The quantitative data were analyzed by unpaired Student t-test and presented as mean ± SEM. n=51. **, P<0.01; ns, no significance. **I.** The proposed mechanism by which MNP33 reprograms cellular bioenergetics and lipid metabolism. The novel microprotein MNP33, originating from an sORF within the Cdhr4 locus, localizes to the inner mitochondrial membrane (IMM) where it interacts with ANT2. This interaction initiates a reprogramming of mitochondrial function: MNP33 modulates ANT2 activity, enhancing the import of glycolytically-derived ATP, which contributes to maintaining a paradoxically high membrane potential (ΔΨm) despite MNP33 also causing increased proton leak. This confluence establishes a “wasteful” bioenergetic state characterized by inefficient ATP production via oxidative phosphorylation. Concurrently, MNP33 reprograms cellular lipid metabolism: it decreases fatty acid uptake and alters intracellular fatty acid partitioning, reducing incorporation into triacylglycerols (TAG) while increasing incorporation into cholesteryl esters (CE). This shift towards CE storage is also linked to MNP33-mediated stabilization of the key enzyme ACAT1, potentially facilitated by closer ER-mitochondria contacts. Downstream cellular consequences include increased ROS production driving autophagy/mitophagy, and inhibited cell proliferation without inducing apoptosis. The illustration was created with BioRender.com.

To further understand how MNP33 promotes CE accumulation, we focused on ACAT1, the key enzyme responsible for converting free cholesterol to CE. Western blotting analysis of subcellular fractions revealed a significant increase in ACAT1 levels in the mitochondrial and total membrane fractions of MNP33-overexpressing cells treated with or without OA, whereas no change was observed in the post-nuclear supernatant (PNS) fraction (Figure 6E). HMGCR, the rate-limiting enzyme in cholesterol synthesis, was not affected by MNP33 overexpression in all subcellular fractions.

Further investigation using the proteasome inhibitor MG132 suggested that MNP33 stabilizes ACAT1 in mitochondria. In control cells, MG132 treatment significantly increased ACAT1 levels in isolated mitochondria (Figure 6F, lane 3 vs lane 1). However, in MNP33-overexpressing cells, the MG132-induced increase in mitochondrial ACAT1 was less pronounced (Figure 6F, lane 4 vs lane 2), suggesting that MNP33 already stabilizes ACAT1, reducing its degradation. ACAT1 localizes to the ER and mitochondrion-associated membranes (MAMs)[34]. TEM analysis revealed that MNP33 overexpression significantly shortened the distance between ER and mitochondria, although the contact length remained unchanged (Figure 6G and 6H).

These findings suggest that MNP33 may stabilize ACAT1, potentially facilitated by the observed closer interactions between ER and mitochondria, thereby enhancing CE accumulation.

## Discussion

Recent research has uncovered a multitude of previously overlooked “dark proteins” produced from regions of the genome not traditionally considered protein-coding. These often small and unstable proteins are being studied for their potential roles in cellular functions. Previously, we established a novel system to identify potential proteins derived from noncoding RNAs on lipid droplets, focusing initially on isolated lipid droplets (LDs) and using mass spectrometry against a dedicated sORF-derived protein database. This organelle-focused “bottom-up” strategy differs from broader approaches such as ribosome profiling or whole-cell proteomics by incorporating enrichment steps for small proteins directly from organelles, thereby improving the detection of low-abundance or unstable microproteins. Using this system, we previously identified 44 candidates, including LDANP1 on LDs[24, 25]. The approach may be extended to identify microproteins localized to other organelles.

Intriguingly, among the 44 proteins identified from the LD fraction, multiple candidates, including MNP33 characterized here, were found to localize to mitochondria. This might seem counterintuitive but is consistent with known close interactions between LDs and mitochondria— particularly in metabolically active tissues such as skeletal muscle, heart, and brown adipose tissue— where these associations facilitate lipid transfer to meet energy demands[35, 36]. Proteomic studies of isolated LDs from those tissues or cells often co-identify numerous mitochondrial proteins, suggesting either proteins involved in direct LD-mitochondrion contacts or the presence of distinct mitochondria subpopulations tightly associated with LDs[35, 37, 38]. Thus, our LD-focused screen likely captured MNP33 due to its presence in mitochondria closely associated with LDs. We suggest that both LD protein co-immunoprecipitation[39], using LD-resident proteins such as ADRP/PLIN2 or TIP47/PLIN3, and the adiposome binding assay[40] could serve as effective approaches for identification of LD-associated dark proteins in future studies.

This study provides the first in-depth functional characterization of MNP33, a novel 28-amino acid protein confirmed here to reside in the inner mitochondrial membrane and originating from a transcript region within the *Cdhr4* gene locus previously considered noncoding. Its discovery adds to the growing functional importance attributed to the “dark proteome”. While detecting endogenous MNP33 protein proved challenging, likely due to low abundance or rapid turnover, several lines of evidence support its genuine expression and translation. MNP33 is highly conserved among mammals (Figure 2A), suggesting functional constraints. Furthermore, analysis of published mouse testis ribosome profiling data confirms ribosome occupancy over the *Mnp33* sORF with high significance, leading to its annotation SPROMMU107011 in the SmProt database (http://bigdata.ibp.ac.cn/SmProt/Contact.htm) (Figure S4A and S4B)[41, 42]. Additionally, the MS2 spectrum used for its initial identification is of high quality and matches the calculated molecular weight, collectively supporting endogenous MNP33 expression (Figure S4C).

Our functional investigation in this study revealed that MNP33 profoundly impacts mitochondrial structure and function. Its overexpression induced significant mitochondrial elongation and fostered a paradoxical bioenergetic state: respiration became less coupled, exhibiting increased proton leak, yet ΔΨm was maintained at a high level. This points towards a fundamental reprogramming of mitochondrial priorities. The interaction between MNP33 and ANT2 appears pivotal in rewriting these bioenergetics. Our data indicate a strong interaction and suggest MNP33 modulates ANT2 activity, supporting a model where MNP33 enhances ANT2’s electrogenic ATP/ADP exchange, fueled by increased glycolysis, to bolster ΔΨm. This mechanism allows mitochondria to sustain high potential even under stress, suggesting a prioritization of potential maintenance over maximal ATP yield efficiency.

Integrating these findings with known ANT functions suggests the MNP33-ANT2 interaction acts as a critical regulatory node with potentially broader consequences. Beyond bolstering ΔΨm, it is plausible, given reports linking ANT proteins to proton conductance[43], that MNP33’s modulation of ANT2 also directly contributes to the increased proton leak. This would mechanistically link the interaction to the observed decrease in ATP production efficiency. Furthermore, considering links between ANT dysfunction and ROS generation[44], the MNP33-ANT2 interaction could also plausibly influence the increased ROS levels, perhaps directly or indirectly. Thus, the MNP33-ANT2 interaction may represent a central mechanism simultaneously impacting potential, coupling efficiency, and oxidative stress.

This altered mitochondrial state triggered significant cellular stress responses. MNP33 overexpression led to increased ROS production, likely contributing to the observed induction of autophagy/mitophagy, while markedly inhibiting cell proliferation. Notably, apoptosis was not induced, suggesting activation of adaptive survival pathways over cell death.

Concurrently, MNP33 expression remodels lipid metabolism, favoring cholesteryl ester storage over triacylglycerols. This correlates with increased stability of ACAT1, the key CE synthesis enzyme. How does MNP33 achieve this? Our primary hypothesis links this stabilization to the observed structural changes at the ER-mitochondria interface. MNP33 expression significantly shortens the distance between these organelles (Figure 6G), and we propose that these closer MAM contacts create a microenvironment favoring ACAT1 stability. While this provides a plausible explanation linking the structural and biochemical changes, further studies are needed to confirm causality, as other mechanisms stabilizing ACAT1 may also be involved. A potential mechanism underlying MNP33-mediated reprogramming of cellular bioenergetics and lipid metabolism is depicted in Figure 6I.

In conclusion, this study identifies MNP33 as a novel, functionally significant inner mitochondrial membrane protein encoded by a supposedly non-coding region. MNP33 acts as a potent regulator of mitochondrial function and cellular metabolism, likely exerting many of its effects via interaction with ANT2, leading to altered bioenergetics, lipid handling, and stress responses. Our work highlights the functional importance hidden within the “dark proteome” and presents MNP33 as a potential target for modulating mitochondrial function and combating metabolic disease at the cellular level. Future studies should aim to elucidate the precise molecular mechanisms of MNP33’s interactions and the full scope of its regulatory network within the cell.

## Acknowledgments

We thank Liqing Liu at the Center for Biological Imaging (CBI), Institute of Biophysics, Chinese Academy of Sciences, along with Zihan Chen and Li Xiao from Beijing University of Posts and Telecommunications, for performing the DeepContact analysis.

This work was supported by the National Key R&D Program of China (Grant No. 2023YFA1801103 and 2024YFA1306101), National Natural Science Foundation of China (Grant No. 92357302 and 32170787) and Beijing Research Ward Excellence Program (Grant No. BRWEP2024W102170106).

## Author contributions

S.Z., P.L. and T.L. conceived and designed the project. Y.X., Q.Y., Y.W., Q.L., X.F., Z.C., B.P., K.Z, and J.W. performed the experiments. S.Z., P.L. and Y.X. wrote the manuscript. All authors have read and approved the manuscript.

